# Endogenous sugar level is associated with differential heat tolerance in onion bulb scales

**DOI:** 10.1101/848572

**Authors:** Ortal Galsurker, Gilor Kelly, Adi Doron-Faigenboim, Kalaivani Aruchamy, Bolaji Babajide Salam, Paula Teper-Bamnolker, Amnon Lers, Dani Eshel

## Abstract

Postharvest heat treatment stimulates desiccation and browning of outer scales of onion (*Allium cepa*. L) bulb to dry papery skins. Inner scales resist the heat treatment, as evidenced by high moisture levels. During heating, inner scales showed increasing soluble sugar levels followed by higher osmolarity, vs. a dramatic decrease in both in the outer scales. Exogenous feeding of outer scales with sucrose, glucose or fructose solutions before heat treatment reduced water loss during heating, suggesting a role for soluble sugars in water retention and therefore, heat tolerance. Vacuolar invertase (VInv) is a key enzyme regulating the levels of sucrose, glucose and fructose in plant tissue. *VInv*-silencing in potato plants prevented the accumulation of reducing sugars in heated leaves, increasing water loss. In onion outer scales, VInv activity increased during heating but reducing sugars decreased, possibly due to their rapid metabolism during scale senescence to form skin. Transcriptomic analysis demonstrated upregulation of genes involved in lignin biosynthesis and secondary cell-wall formation in outer scales during heat exposure, and upregulation of genes involved in energy-related pathways in inner scales. This study reveals the dual role of soluble sugars in different onion scales, as osmoprotectants or building blocks, under heat stress.

**Highlight:** Hexose formation in the inner scales of onion is associated with heat tolerance, while in the outer scales, the hexoses are metabolized for onion skin formation.

## Introduction

Onion (*Allium cepa* L.), a widespread Alliaceae plant, is one of the main vegetables consumed worldwide. Hot-air curing of bulbs is an important postharvest treatment, used to dry out the outer scales, which are then transformed into a complete skin. This skin protects the bulb from water loss and suppresses disease incidence, thereby maintaining higher quality during storage (Chope *et al*., 2012; Downes *et al*., 2009; Maw *et al*., 2004). Application of postharvest heat treatment, 33 °C at 98% relative humidity (RH) for a few days to a few weeks, to detached outer and inner scales reveals their differential responses to the heat stress (Galsurker *et al*., 2018). The outer scale desiccates and turns into papery dry skin, while the inner scale exhibits tolerance to the heat stress, maintaining high relative water content (RWC; Galsurker *et al*., 2018). The mechanism responsible for these scales’ differential heat response is unknown.

Plant tissue copes with heat stress through a series of biochemical and metabolic changes; among these is the accumulation of compatible solutes that help the plant reestablish osmotic homeostasis by increasing the water potential, protecting cellular organelles and stabilizing proteins and membranes (Hasanuzzaman *et al*., 2013). The accumulated compatible solutes, also termed osmoprotectants, are low-molecular-weight organic solutes that are highly soluble and do not affect plant metabolism (Yancey, 2005). Several soluble sugars are considered osmoprotectants, shown to accumulate in plants in response to abiotic stresses, including high temperature, and to maintain cell homeostasis (Gepstein *et al*., 2008; Hare *et al*., 1998; Vinocur and Altman, 2005).

Onion bulbs contain fructose, glucose, sucrose and a series of fructo-oligosaccharides (fructan) as the main non-structural carbohydrates, accounting for 80% of bulb dry matter (DM; Benkeblia *et al*., 2004; Benkeblia *et al*., 2002; Darbyshire and Henry, 1981; Darbyshire and Henry, 1979). During bulb storage, metabolic activities lead to quantitative variations in sugar composition which are strongly related to the transition from dormancy to sprouting (Benkeblia and Varoquaux, 2003; Benkeblia *et al*., 2002). However, there are conflicting reports on the nature of the changes in soluble sugars in onion bulbs during storage. After prolonged storage, a considerable increase in the amount of fructose and glucose was reported (Chope *et al*., 2007). In other studies, fructose levels were also reported to increase during the first month of storage but then they decreased (Benkeblia and Varoquaux, 2003; Benkeblia *et al*., 2002). In contrast, Downes *et al*. (2010) reported a decrease in fructose level during initial storage followed by a sudden increase after 12 weeks of storage. Although there are many studies describing the sugar alterations during onion bulb storage, the nature of the changes in sugar content is not yet clear and there are no reports evaluating the soluble sugar levels and their functions in different onion scales in the same bulb.

The main aims of this study were to assess the role of soluble sugars in the differential heat response of outer and inner onion bulb scales. We showed that high soluble sugars promote heat tolerance in the inner scale. In the outer scale, on the other hand, we found that sugars are rapidly metabolized to form skin tissue.

## Materials and methods

### Plant materials and heat treatment

Commercial brown onion cv. Orlando was grown in sandy soil in the northwestern Negev desert, Israel, in the years 2015–2017. The onions were not treated with maleic hydrazide before leaf drop, the common agricultural practice, and did not undergo field curing. Onions were harvested manually at 80–100% fallen leaves (top–down) and the leaves were removed with a sharp knife, leaving a ~5-cm long neck above the bulb, as described previously (Eshel *et al*., 2014). Dry muddy skin was removed to expose the first scale, and bulbs were separated into different successive scales, which were numbered from the exterior to interior of the bulb as described previously (Galsurker *et al*., 2017). The first scale represents the first outer scale which has the ability to form additional skin following heat treatment, and the fifth scale (outside in) represents an inner fleshy scale.

Heat treatment for the detached outer and inner scales was performed as described by Galsurker (2018). Detached scales were incubated at 33 °C under 98% RH in an incubator (Binder KBF720, Binder Instrument Co., Germany) and sampled over time. Powdered scale material was prepared by freezing the individual scales in liquid nitrogen and grinding to a fine powder using a liquid nitrogen grinder/mill (IKA, Germany). The freeze-dried powdered scales were kept at −20 °C until use.

Since DNA transformation is still a challenge in onion, transgenic potato leaves were used to demonstrate the wider perspective of our findings. Leaves of potato (*Solanum tuberosum* ‘Russet Burbank’ [RBK]) were harvested from previously developed vacuolar invertase (*VInv*)-silenced lines (RBK1, RBK22 and RBK46; Zhu *et al*., 2014). The fourth true leaves of same-age plants were detached simultaneously, transferred to heat treatment at 30 ^o^C and 65% RH, and sampled over 6 days.

### Osmolarity measurements

To measure onion scale osmolarity, 1-cm diameter discs were punched from the first (outer) and fifth (inner) scale with a cork borer. Tissue discs were immediately frozen in liquid nitrogen and then allowed to thaw for 30 min at room temperature (RT). Thawed scales were ground and extracts were centrifuged at 13,000 *g* for 10 min at 4 ºC; the scale cell sap was collected. The collected sap (10 µL) was used to determine osmolalrity using a vapor-pressure osmometer (Vapro 5500, Wescor, USA) according to the manufacturer’s instructions. To measure potato leaf osmolarity, 0.5 g of leaf was placed in a 50-mL tube, then immediately dropped into liquid nitrogen and stored at −80 °C until sap extraction. Frozen leaves were loaded into a 10-mL syringe covered with a filter‐paper disc and thawed at RT. Thawed leaves were centrifuged at 1,500 *g* for 10 min at 4 ºC and leaf sap was collected to determine the osmolarity as described above.

### Extraction and quantification of sugars

Powdered scale material (150 mg DW) and potato leaves (0.5 g FW) were heated three times with 80% ethanol at 80 °C, for 45 min each time, and the ethanolic solutions were pooled and dried using a speed vacuum (Centrivap Concentrator, Labconco, USA). Pelet was dissolved in 2 ml DDW and passed through a 0.2-μm membrane filter (Millex-GV Filter Unit, Merck Millipore, Ireland). The filtrate was used for sucrose, glucose and fructose analysis by ultrafast LC (LC-10A UFLC series, Shimadzu, Japan) equipped with a refractive index detector (SPD-20A) and analytical ion-exchange column (6.5 x 300 nm; Sugar-Pak I, Waters). Column temperature was set to 80 °C and ultrapure water (Bio Lab, Israel) was used as the mobile phase, at a flow rate of 0.5 mL min^−1^. Sugars were identified and quantified by a calibration curve created using standards of sucrose, glucose and fructose. The chromatographic peak corresponding to each sugar was identified by comparing the retention time with that of the standard. The calibration curve was used to determine the relationship between the peak area and concentration of the sugars.

### Exogenous feeding of sugars to outer scales

Detached outer scales were surface-sterilized with 0.1% (v/v) sodium hypochlorite and 0.02% (v/v) Tween 20 for 5 min, followed by washing with sterile water for 10 min. The outer scales were then dried for 5 min and placed in a 500-mL sterile beaker containing 0, 100, 300 or 600 mM sucrose, glucose or fructose solutions, and incubated at 14 °C in the dark for 24 h. After incubation, outer scales were carefully rinsed with sterile water for 5 min and exposed to heat treatment as described above, for 6 days. Scales were sampled daily during the heating for osmolarity and water-loss measurements.

### Relative water content

Water loss from the exposed scale was determined by measuring RWC during the heat treatment, as described by Galsurker (2018). The heated detached scales were sampled as follows: six 2-cm diameter discs were punched from each scale with a cork borer and weighed for initial FW. The discs were then soaked in distilled water for 24 h at RT, and carefully blotted dry with tissue paper to determine their saturated weight (SW). The discs were then dried in an oven at 70 °C for at least 24 h to measure the DW. RWC was calculated as:

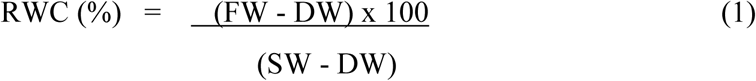

### Enzyme extraction and activity measurements

VInv activity was measured as described previously (Miron and Schaffer, 1991), with minor modifications. Outer and inner powdered scale material (1 g DW) was dissolved in 5 mL extraction buffer containing 25 mM HEPES-NaOH, 7 mM MgCl_2_, 0.5 mM EDTA, 3 mM DTT, and 2 mM diethyldithiocarbamic acid, pH 7.5. After centrifugation at 18,000 *g* for 30 min, the supernatant was dialyzed overnight against 25 mM HEPES-NaOH and 0.25 mM EDTA, pH 7.5, and used as a crude extract. VInv activity was measured by incubation of 0.3 mL of 0.1 M citrate/phosphate buffer (pH 5.0), 0.1 mL crude extract and 0.1 mL of 0.1 M sucrose. After 1 h incubation at 37 ^o^C, glucose liberated from the hydrolysis of sucrose was quantified by adding 500 μL Sumner’s reagent (3,5-dinitrosalicylic acid) and immediately transferring the sample to heating at 100 ^o^C for 10 min to terminate the reaction. The sample was then chilled at 4 °C (Sumner and Graham, 1921). The reduction of dinitrosalicylic acid to 3-amino-5-nitrosalicylic acid by glucose was measured as absorbance at 550 nm using a spectrophotometer. Quantitation of glucose in each sample was based on glucose standards. VInv activity was expressed as nmol glucose formed g DW^−1^ min^−1^.

## Results

### Heat treatment induces different osmolarity profiles in outer vs. inner scales

Our previous study showed that the first outer onion scale dramatically desiccates via loss of RWC during heat treatment, while the fifth inner scales maintain their RWC and are thus termed heat tolerant (Galsurker *et al*., 2018). To determine the factors that might be involved in the differential heat response, detached outer and inner onion scales were heated at 33 °C and 98% RH and their osmolarity measured over 6 days. Differential osmolarity levels were found between outer and inner scales during the treatment. The inner scale revealed a high osmolarity level that increased slightly from 455 to 528 mosmol kg^−1^ during the first 2 days of heating and then stabilized until day 6 (Fig. 1A). In contrast, the osmolarity of the outer scale, which initially had lower levels of 409 mosmol kg^−1^, decreased gradually to 276 mosmol kg^−1^ during the 6 days of heat treatment, suggesting that outer scale desiccation is associated with a decline in osmolarity level (Fig. 1A).

**Fig. 1.**
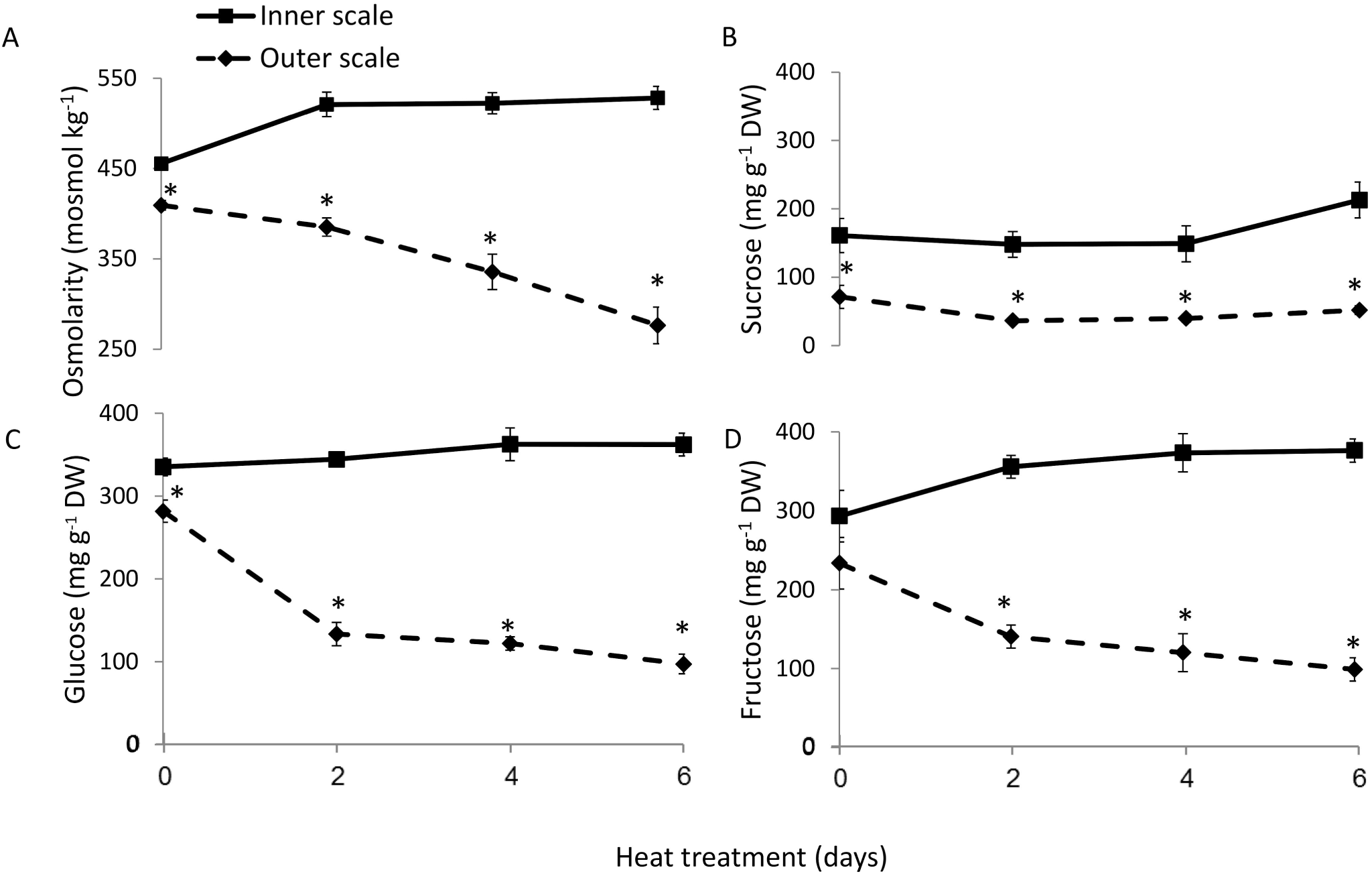
Heat treatment induces differential changes in osmolarity and soluble sugar levels in the outer vs. inner scales of onion bulb. Detached onion scales were exposed to 6 days of heat treatment (33 °C, 98% RH). Data are averages of three experiments, each performed with five replicates per treatment. Error bars represent SE (n = 6). A single asterisk (*) represent significant differences (P < 0.05) between different onion scales among the same time point.

### The inner scale maintains high levels of soluble sugars under heat treatment

Our previous data suggested that heat treatment of outer and inner scales induces the differential expression of various genes, including those related to sugar metabolism (Galsurker *et al*., 2018). To test whether soluble sugars contribute to the different osmolarity levels observed between the inner and outer scales, we quantified sucrose, glucose and fructose levels during the heat treatment. Significant differences in total soluble sugars between inner and outer scales were already observed at the 0 time point, which became more significant following 2, 4 and 6 days of heating. The inner heated scale showed higher initial levels of glucose, fructose and sucrose and a slight increase in the levels of these sugars during the 6 days of heating (Fig. 1B–D). The outer scale showed a decrease in all three soluble sugars’ levels, especially in the first 2 days of heating (Fig. 1B–D). The contents of the hexoses glucose and fructose decreased dramatically in the outer scale (Fig. 1C, D). The association between the soluble sugar and osmolarity profiles of the onion scales and the desiccation of the outer scale suggested a role for sugar metabolism in heat susceptibility.

### Exogenous sugar feeding induces higher osmolarity levels and heat tolerance

We hypothesized that soluble sugar content directly contributes to the observed increase in osmolarity and reduced water loss in the inner scale tissue during exposure to heat. We tested this hypothesis by feeding outer scales with a solution containing glucose, fructose, or sucrose and followed the consequences on osmolarity and water loss. Detached outer scales were immersed in 100, 300 and 600 mM solutions of glucose, fructose or sucrose, and incubated in the dark at 14 °C for 24 h. To detect sugar penetration into the scale tissue, we quantified its level after sugar feeding, prior to the heat treatment. Higher levels of sugars were found in the outer scales that were fed with sugars compared to the non-fed controls, demonstrating that the three sugars could penetrate the scale (Supplementary Fig. S1). After sugar feeding, the scales were exposed to heat treatment for 6 days. Glucose, fructose and sucrose feeding only increased osmolarity at the higher doses of 300 and 600 mM, with no significant differences among sugars (Supplementary Fig. S2). During heating, the scales that were fed sugars maintained high osmolarity levels, while osmolarity decreased in the non-fed scales (Fig. 2A). Scales that were fed with any of the three sugars retained high RWC values, between 85 and 90%, compared to the non-fed scales that showed an average 82% RWC after 6 days of heat treatment (Fig. 2B). This experiment suggested that elevated sugar content reduces outer scale desiccation through an increase in tissue osmolarity.

**Fig. 2.**
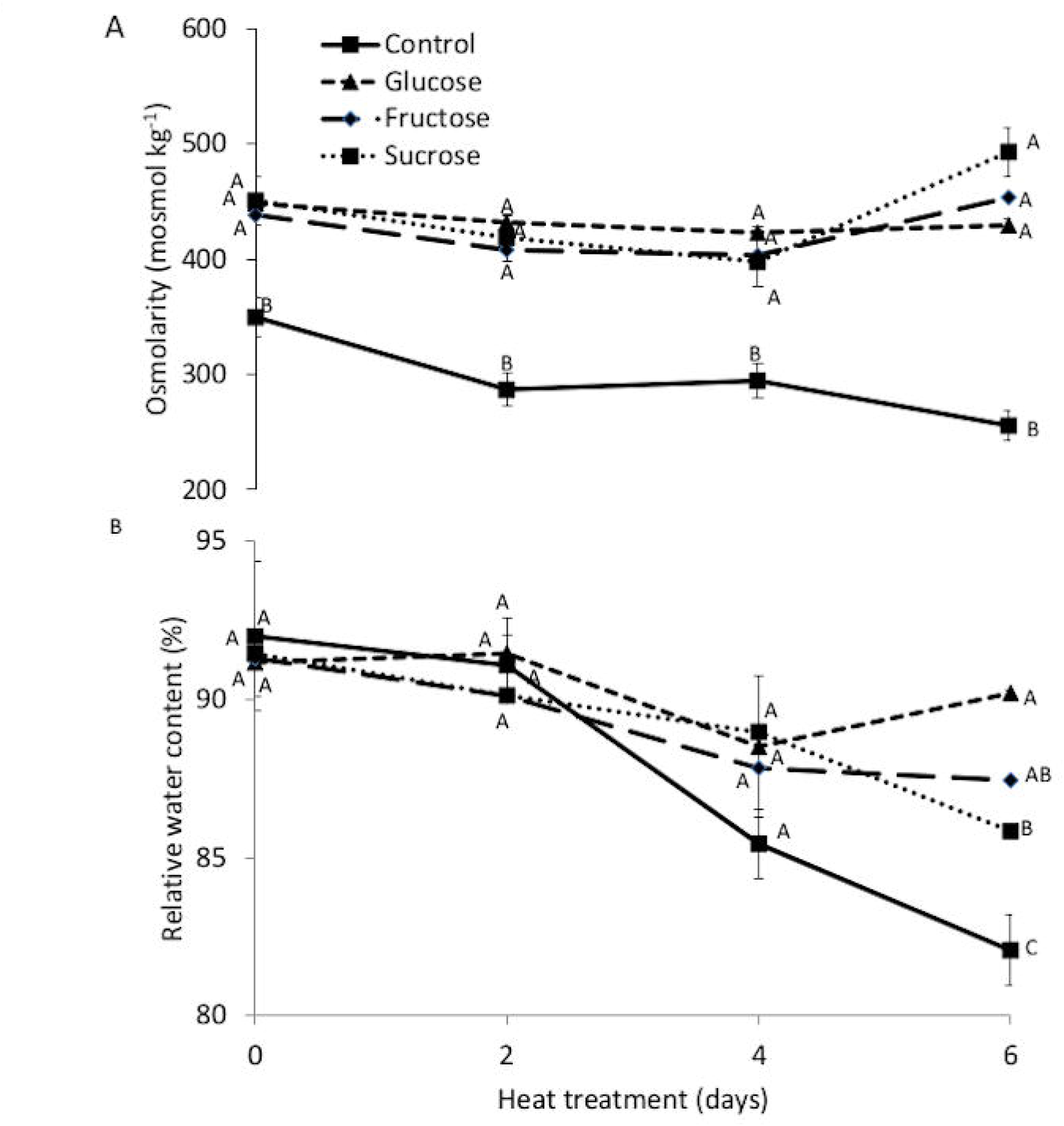
Sugar feeding stabilizes tissue osmolarity and decreases water loss during heat treatment. (A) Osmolarity and (B) relative water content of detached onion scales during 6 days of heat treatment (33 °C, 98% RH). Outer scales were incubated in 600 mM sucrose, glucose or fructose solutions for 24 h at 14 °C, before the heat treatment. Data are averages of three experiments, each performed with five replicates per treatment. Error bars represent SE. Different uppercase letters indicate significant statistical difference (P < 0.05) between different sugars among the same time point.

### *VInv* silencing increases susceptibility to heat

VInv is a key enzyme responsible for the level of sucrose and its hydrolysis products, glucose and fructose. Silencing of *VInv* in potato plants results in effective prevention of sucrose degradation (Bhaskar *et al*., 2010; Salam *et al*., 2017). To study the possible involvement of osmolarity, as affected by soluble sugar content, on heat tolerance of plant tissue, we used three independent *VInv*-silenced lines of RBK potato (Zhu *et al*., 2016; Zhu *et al*., 2014). Leaves from these lines have reduced levels of glucose and fructose (Zhu *et al*., 2014). During 8 days of heat treatment (30 ^o^C at 65% RH), leaves of the three *VInv*-silenced lines— RBK1, RBK22 and RBK46—lost more water than the wild-type (WT) leaves (Fig. 3). The levels of all three soluble sugars decreased in the three *VInv*-silenced RBK lines during the heat treatment. A different sugar profile was found in the WT, where the sucrose levels decreased and the hexose levels increased during heating (Fig. 3B). This result fit with the expected activity of VInv that cleaves sucrose and increases the level of hexoses in association with higher tolerance to heat.

**Fig. 3.**
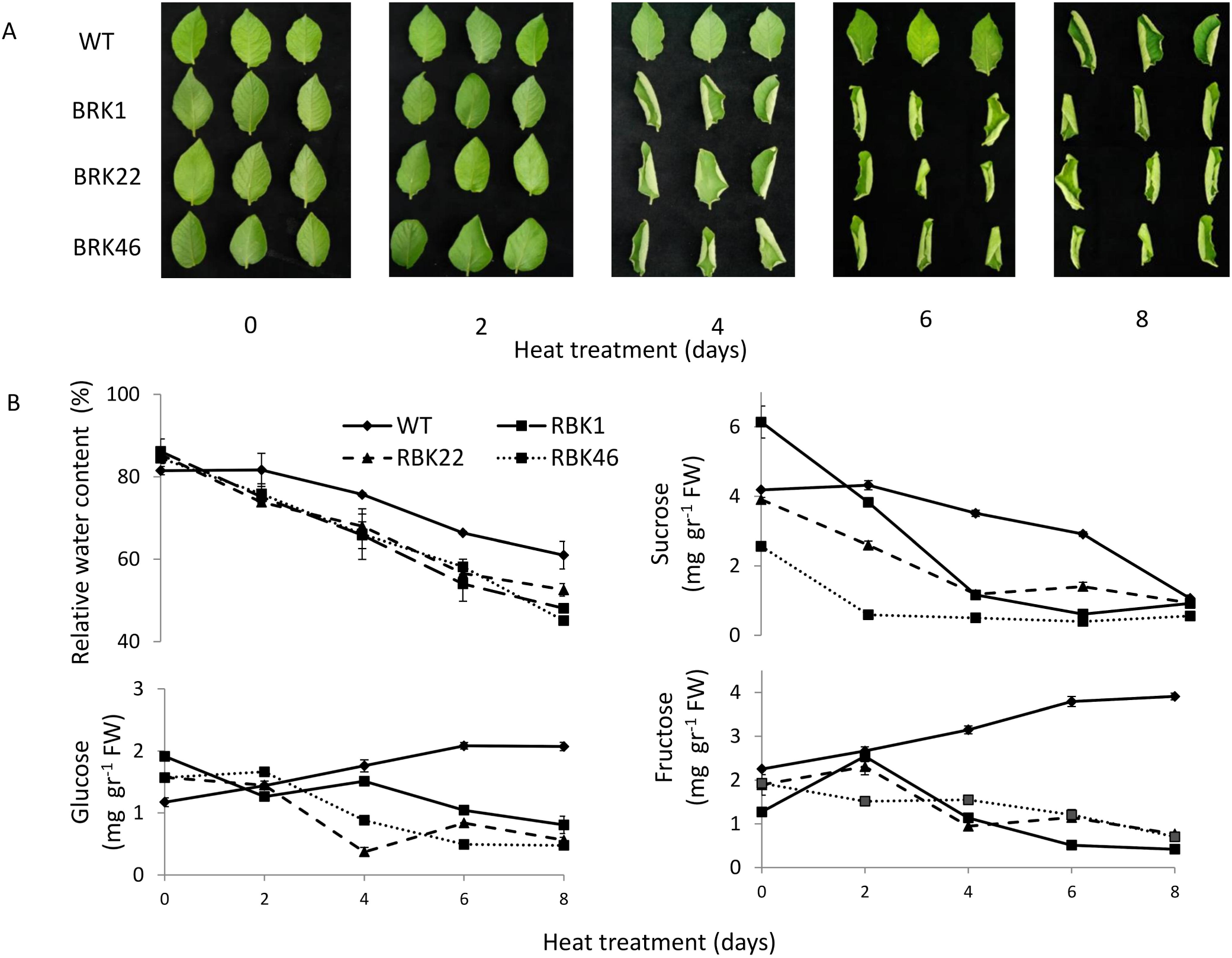
Silencing *VInv* enhances desiccation of potato leaves during heat treatment. Leaves of RBK1, RBK27 and RBK46—*VInv*-silenced lines of potato cv. Russet Burbank—and wild type (WT) were exposed to 8 days of heat treatment (30 ^o^C, 65% RH). (A) Phenotypic documentation, every 2 days, of 3 leaves from 3 plants representing each line. (B) Relative water content. Eror bars represent ± SE of three repeats, each with 10 leaves.

### Heat treatment induces accumulation of VInv activity only in the outer scale

We hypothesized that sugar metabolism in the onion scale contributes to the increase in osmolarity, similar to the effect of sugar feeding (shown in Fig. 2). As supported by the consequences of *VInv* silencing in potato leaves, this enzyme may play a role in determining sugar levels and consequently, water loss during heat stress. We examined possible VInv involvement in the heat response in onion as well, by measuring the changes in enzyme activity during heat treatment in the outer and inner scales. Surprisingly, after 8 days of heat treatment, VInv activity had increased dramatically in the outer scale, whereas in the inner scale, it remained constant (Fig. 4A). The two peaks of VInv activity observed in the outer scale during the heat treatment (Fig. 4A) suggested a partial inhibitory effect caused by the accumulation of hexoses, as suggested by Kim *et al*. (2000). However, these results did not fit with the measured changes in soluble sugar levels in the outer and inner scales during the heat treatment (Fig. 1). One possibility for this discrepancy is that the reduction in hexose content observed in the outer scales, even as VInv activity increases, is related to hexose recruitment for DM synthesis in the drying outer scale tissue.

**Fig. 4.**
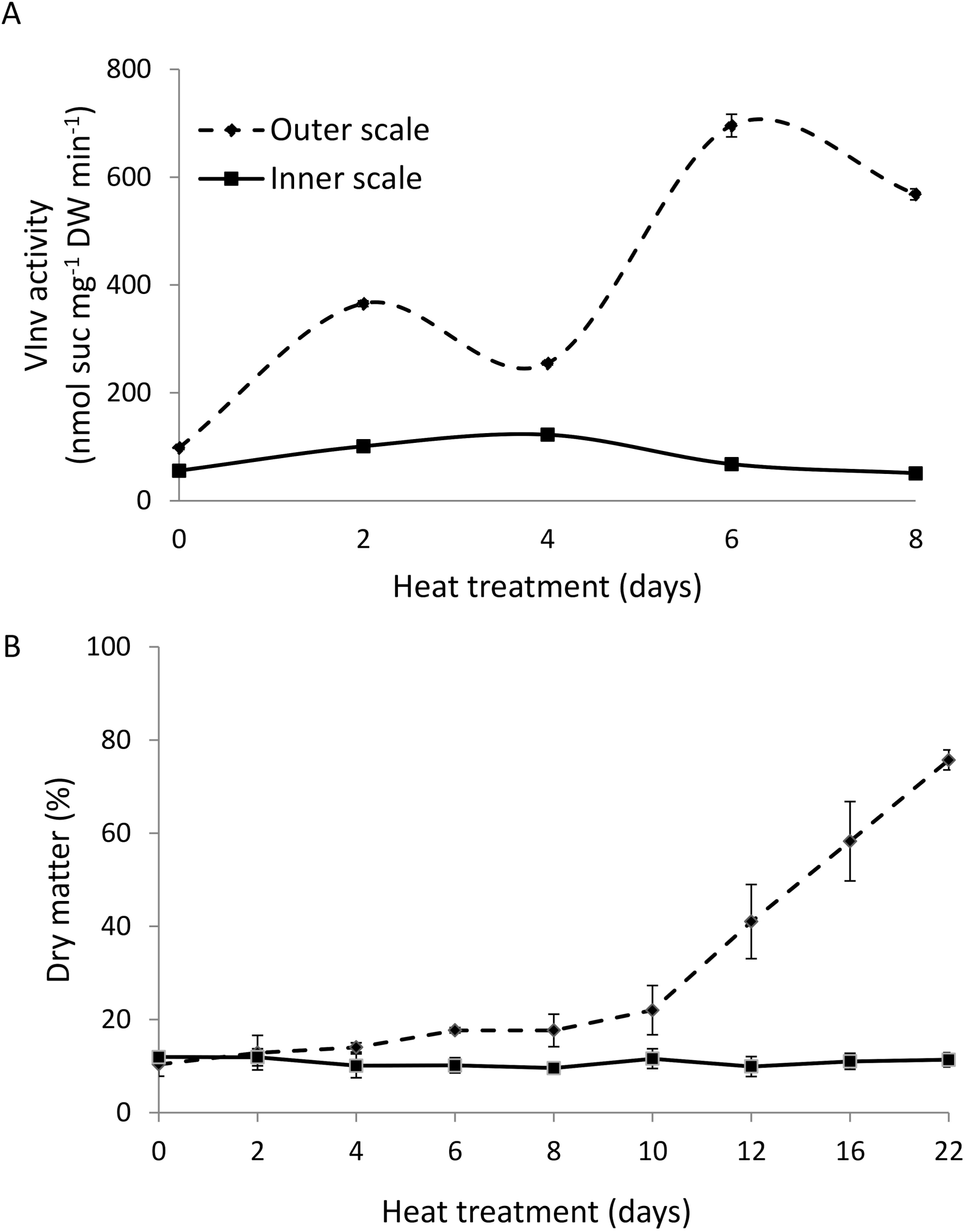
Heat treatment induces accumulation of (A) *VInv* activity and (B) DM only in the outer scale of the onion bulb during 8 and 22 days of heat treatment (33 °C, 98% RH), respectively. Data are averages of three experiments, each performed with five replicates per treatment. Error bars represent SE.

To examine this possibility, DM content was compared between the outer and inner scales during prolonged heat treatment, and the profiles were markedly different. In the outer scale, a gradual increase in DM contents, from 11% to 75%, was measured during 22 days of heating, which eventually resulted in skin formation, whereas the inner scales maintained a constant DM content of about 11% (Fig. 4B). The increase in DM content in the outer scale was probably a result of higher cell-wall development during scale desiccation and skin formation. Cellulose and lignin are two of the major components of the secondary cell wall, serving as barriers against pathogen colonization of plant tissue (Miedes *et al*., 2014). We reasoned that upregulation of genes related to the biosynthesis of cellulose and lignin would be positively correlated with skin formation in the outer scale and the observed increase in DM.

Analysis of our heat treatment transcriptome data (Galsurker *et al*., 2018) showed upregulation of several genes involved in lignin biosynthesis and secondary cell-wall formation in the outer scale (Fig. 5A, B; Supplementary Table S1). These results suggest that the soluble sugars in the outer scales are used as building blocks for the synthesis of structural carbohydrates, and thus their levels decrease as part of outer skin development.

**Fig. 5.**
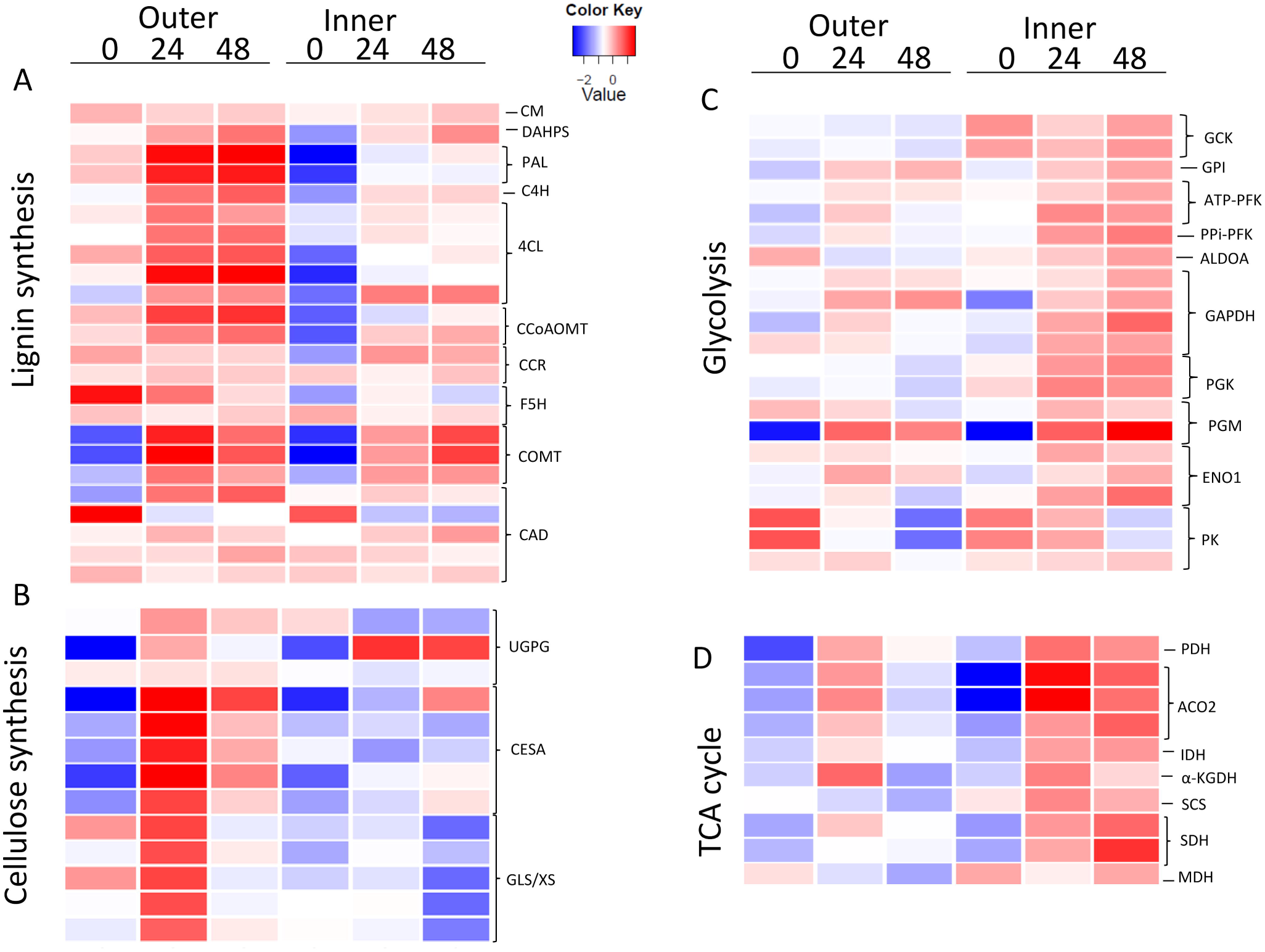
Heat map describing the expression profiles of genes related to sugar metabolism in the outer vs. inner scale. Genes related to (A) lignin synthesis, (B) cellulose synthesis, (C) glycolysis and (D) TCA cycle. Measurements were performed at 0, 24 and 48 h of heat treatment in the outer and inner scales.

The inner scale showed heat resistance and maintenance of viability. Genes involved in energy-related pathways, such as glycolysis and the tricarboxylic acid (TCA) cycle, were upregulated in the inner scale during the heat treatment (Fig. 5C, D). After glycolysis, the respiratory mechanism continues with the TCA-cycle reactions. The pyruvate produced in glycolysis can be transported to the mitochondria where it is oxidized to acetyl-CoA and CO_2_ by pyruvate dehydrogenase (Werner *et al*., 2011). In the inner scale, genes involved in the TCA cycle were significantly induced following the heat treatment, and were highly expressed compared to the levels measured in the outer scale (Fig. 5D).

## Discussion

### A high level of soluble sugars promotes heat tolerance in the inner scale

Only outer scales of onion can form dry brown skin during heat stress, while the inner scales maintain high water content and do not change color (Galsurker *et al*., 2018). During heating, osmolarity and hexose levels were reduced in the outer scale, and were stable in the inner one, suggesting their possible involvement in maintaining the osmotic pressure of internal onion scales (Fig. 1). Such high osmotic pressure could have a role in the heat tolerance of the inner scales. The high level of osmolarity in the inner scale could be achieved through the accumulation and/or maintenance of the three soluble sugars, glucose, fructose and sucrose, during heating (Fig. 1). Soluble sugar accumulation has been confirmed to play an important role in enhancing plant tolerance to heat stress (Hasanuzzaman *et al*., 2013). Soluble sugars such as glucose, sucrose, fructose and trehalose function as osmoprotectants by regulating the osmotic adjustment, provide membrane protection, and scavenge the toxic reactive oxygen species (ROS) formed under various kinds of stresses (Keunen *et al*., 2013; Singh *et al*., 2015). Furthermore, soluble sugars have been reported to participate in the reduction of oxidative damage, partly as a result of activation of specific ROS-scavenging systems (Ramel *et al*., 2009).

Exogenous supply of individual soluble sugars—glucose, fructose or sucrose—to the outer scale led to an increase in its osmolarity and therefore, to a reduction in water loss following heat treatment (Fig. 2), supporting the role of these soluble sugars in osmotic adjustment and heat tolerance. In other studies, exogenous application of glucose during salt stress of wheat seedlings increased DW, maintained ionic homeostasis, induced proline accumulation, prevented water loss and activated antioxidant enzymes (Hu *et al*., 2012). In rice, glucose and fructose functioned as osmoprotectants and free-radical scavengers under salinity stress (Pattanagul and Thitisaksakul, 2008). Furthermore, soluble sugars are associated with ROS anabolism and catabolism, such as the oxidative pentose phosphate pathway involved in ROS scavenging (reviewed by Couée *et al*., 2006). In higher plants, a diverse variety of soluble sugars, such as glucose, sucrose, fructose, raffinose and stachyose are known to provide freezing tolerance (Yuanyuan *et al*., 2009). These sugars not only act as osmoprotectants but also protect membranes by allowing adaptation to drought or chilling stress through their interaction with the lipid bilayer (Garg *et al*., 2002). Our experiments using *VInv*-silenced potato plants showed a decline in reducing sugar content compared to the WT, which increased their susceptibility to heat stress and elevated water loss in their leaves under heat treatment (Fig. 3). These results also support a role for soluble sugars as contributors to osmotic adjustment, thereby promoting heat tolerance in other plant tissues.

### Outer scale sugars are rapidly metabolized during heating

In contrast to the inner scale that maintains high osmolarity, possibly as a result of retaining high levels of soluble sugars, in the outer scale, both osmolarity and hexose levels decreased significantly in response to heat treatment (Fig. 1). Lower levels of hexoses were found to be associated with a higher level of VInv activity, and with increasing DM content in the outer scale under heating (Fig. 4). These results might be explained by rapid metabolism of the hexoses in the outer scale as part of its senescence process. Sugars can act as nutrients as well as regulators of metabolism, growth, stress responses and developmental senescence (O’hara *et al*., 2013; Rolland *et al*., 2006; Rolland *et al*., 2002). The older outer scale, a distressed suicide tissue, is programmed to senesce, die and form skin (Galsurker *et al*., 2017), and all of these process are accelerated by heat treatment. DM content has been previously reported to be correlated with structural carbohydrates in onion scales (Jaime *et al*., 2002). It is possible that sugars in the outer scales are metabolized in processes that result in the synthesis of structural carbohydrates, such as secondary cell-wall components. Various genes associated with secondary cell-wall development were overrepresented in the transcriptome of the outer scale following the heat treatment compared to the inner scales, mainly genes related to lignin and cellulose biosynthesis (Fig. 5). To the best of our knowledge, there are no studies describing the secondary cell-wall development pathway during onion skin formation. However, studies have shown the role of secondary cell-wall development in other protective dry tissues, such as the seed coat. Thickening of the secondary cell wall has been found in seed-coat formation in *Arabidopsis* and *Brassica napus* (Haughn and Chaudhury, 2005; Jiang and Deyholos, 2010).

## Conclusions

The differential response of detached outer and inner bulb scales to heat stress suggests the activation of different cascades of events leading to skin formation and viable heat resistance, respectively (Fig. 6). While the inner scales maintained high osmolarity, DM and sugar levels, all of these factors were dramatically reduced in the outer scales. In both onion scales and potato leaves, high osmolarity, caused by a high level of soluble sugars in the plant tissue, inhibited water loss. The transcriptome analysis in this study led to a putative model for the differential heat responses of the outer and inner scales (Fig. 6). The model suggests that the inner scales express heat tolerance, as compared to the heat susceptibility of the outer scale that results in browning and skin development (Fig. 6). This differential response can be explained by the different physiological ages of the scales, with older ones on the outside and younger ones toward the inside (Brewster, 2008; Galsurker *et al*., 2017). The different biological processes suggested to occur by our study in the inner and outer scales are probably related to their different functions. Outer skins protect the bulb against disease by providing both physical and biochemical barriers to pathogens (Eshel *et al*., 2014; Mishra *et al*., 2014). Upregulation of genes related to lignin and cellulose synthesis in the outer scales, as shown in our transcriptome, represent the cascade that leads to the formation of the biochemical barrier and later, programmed cell death that leads to skin formation (Galsurker *et al*., 2017). In contrast, heat tolerance of the inner scales is associated with higher expression of genes related to glycolysis and the TCA cycle, representing a valuable resource for the identification of candidate heat-tolerant genes, thus providing important information on the mechanisms underlying heat tolerance.

**Fig. 6.**
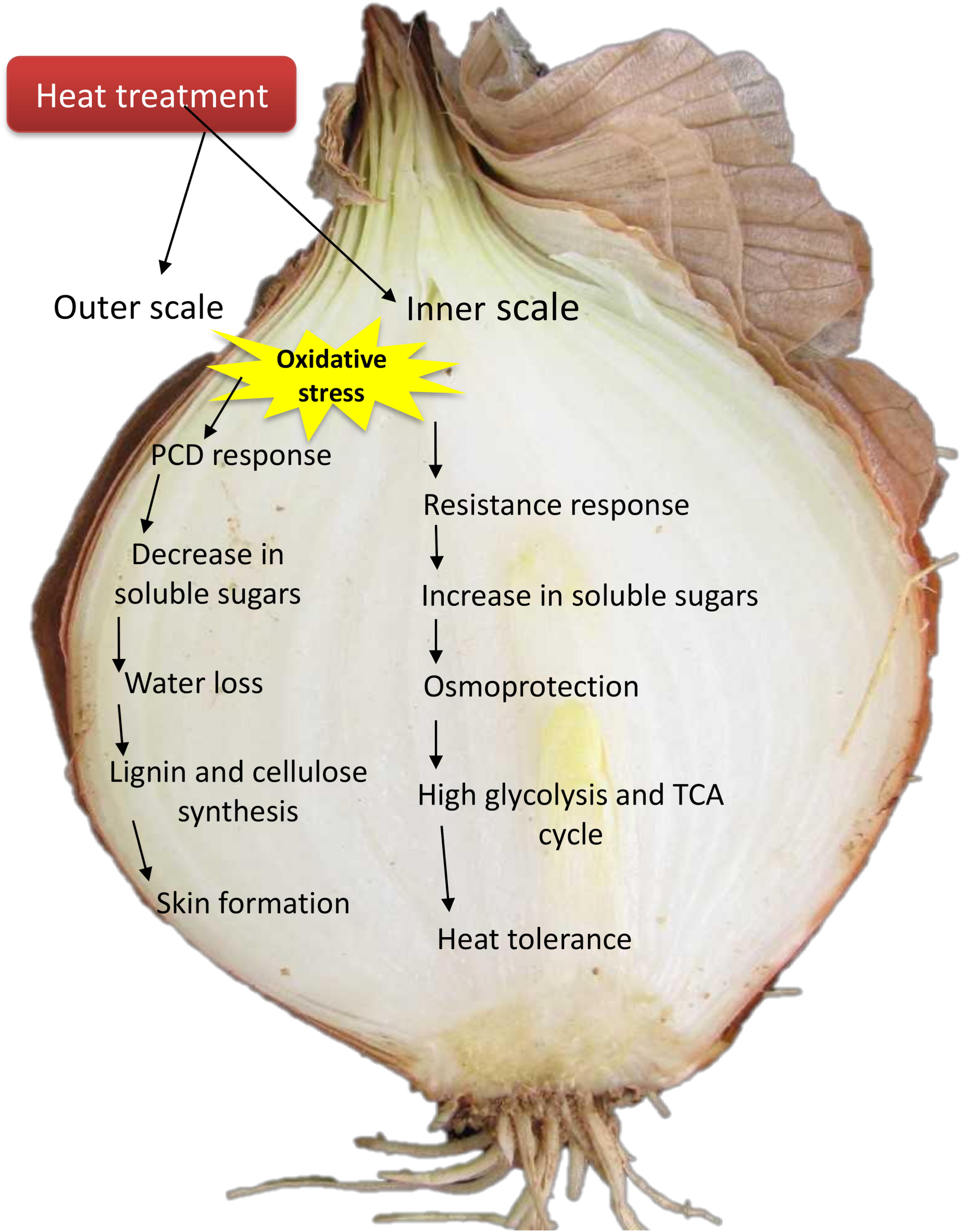
Proposed model for osmoprotection in onion scale in response to heat treatment. The suggested model is based on overrepresentation of the corresponding genes in the first outer and fifth inner scales, as presented in Fig. 5. PCD, programmed cell death.

## Supplementary data

**Table S1.** Gene abbreviations and their full names; presented according to the functional groups in Fig. 5.

**Fig. S1.** Sugar feeding induces a higher level of sugar in onion scale tissue. Outer scales were incubated in 600 mM glucose, fructose or sucrose solution for 24 h at 14 ^o^C. Data are averages of three experiments, each performed with five replicates. Error bars represent SE.

**Fig. S2.** Sugar feeding induces higher osmolarity levels in a dose-dependent manner in the outer scale. Data are means ± SE of six repeats; each repeat contained four detached scales.

## Acknowledgements

This research was funded by a grant from the Chief Scientist of the Ministry of Agriculture and Rural Development of Israel (no. 430-0416-13). We thank Prof. Jiming Jiang for allowing us to work with the RBK transgenic lines. We thank Prof. Menachem Moshelion for fruitful discussions.

